# CRISPR/Cas9-mediated genome editing of *Frankliniella occidentalis,* the western flower thrips, via embryonic microinjection

**DOI:** 10.1101/2023.11.30.569456

**Authors:** Jinlong Han, William Klobasa, Lucas de Oliveira, Dorith Rotenberg, Anna E. Whitfield, Marcé D. Lorenzen

## Abstract

The western flower thrips, *Frankliniella occidentalis*, poses a significant challenge in global agriculture as a notorious pest and a vector of economically significant orthotospoviruses. However, the limited availability of genetic tools for *F. occidentalis* hampers the advancement of functional genomics and the development of innovative pest control strategies. In this study, we present a robust methodology for generating heritable mutations in *F. occidentalis* using the CRISPR/Cas9 genome editing system. Two eye-color genes, *white* (*Fo-w*) and *cinnabar* (*Fo-cn*), frequently used to assess Cas9 function in insects were identified in the *F. occidentalis* genome and targeted for knockout through embryonic microinjection of Cas9 complexed with *Fo-w* or *Fo-cn* specific guide RNAs. Homozygous *Fo-w* and *Fo-cn* knockout lines were established by crossing mutant females and males. The *Fo-w* knockout line revealed an age-dependent modification of eye-color phenotype. Specifically, while young larvae exhibit ivory-colored eyes, the color transitions to bright red as they age. Unexpectedly, loss of *Fo-w* function also altered body color, with *Fo-w* mutants having a lighter colored body than wild type, suggesting a dual role for *Fo-w* in thrips. In contrast, individuals from the *Fo-cn* knockout line consistently displayed bright red eyes throughout all life stages. Molecular analyses validated precise editing of both target genes. This study offers a powerful tool to investigate thrips gene functions and paves the way for the development of genetic technologies for population suppression and/or population replacement as a means of mitigating virus transmission by this vector.

## INTRODUCTION

*Frankliniella occidentalis* (suborder Terebrantia, family Thripidae, subfamily Thripinae), commonly known as the western flower thrips, is a devastatingly invasive crop pest with a global geographical distribution and an extraordinarily broad feeding host range, capable of feeding on hundreds of diverse plant species, tissue types of vegetable crops, ornamentals, tree fruits and fiber crops (reviewed in Reitz et al., 2020). While this thrips species causes direct damage to plants through feeding (piercing-sucking) and oviposition, it is most prominent for its ecological role as a vector of orthotospoviruses (reviewed in Montero-Astúa M. et al., 2016; Oliver & Whitfield, 2016; Rotenberg & Whitfield, 2018), some of the most damaging plant viruses of worldwide economic significance. Management of *F. occidentalis* has been notoriously problematic because it infests both crop and uncultivated plant species, females are highly fecund and populations build up rapidly in a growing season. In addition, grower reliance on broad spectrum chemical control of pest populations has led to thrips resistance to multiple classes of insecticides (Gao et al., 2012) and off-target effects on beneficial arthropods (Bass et al., 2015; Cresswell, 2011; Pisa et al., 2015). Due to the wide-spread importance of *F. occidentalis* in agriculture, it has been a model system for numerous studies centered on pest and vector biology, ecology, insecticide resistance and virus-vector interactions, and it was the first thysanopteran to have its genome sequenced (Rotenberg et al., 2020) and shared with the scientific community (NCBI GenBank accession: GCA_000697945.5).

Although adults of *F. occidentalis* can reproduce sexually, they are haplo-diploid, i.e., haploid males arise from unfertilized eggs (arrhenotokous parthenogenesis), while diploid females develop from fertilized eggs (Moritz, 1997). Adult females oviposit in plant tissue, and depending on food quality and temperature, each female may achieve a reproductive capacity of 200 eggs over its lifespan (Robb & Parrella, 1991). *F. occidentalis* exhibits two wingless larval stages (L1 and L2) that feed on plant leaves and flowering parts. The ensuing immobile and non-feeding pupal stages (propupae, P1; pupae, P2) generally drop to the soil or rest in flowers. Winged adults emerging from the P2 stage exhibit short (Nyasani et al., 2017) or long distance (Mound, 1983) dispersal by flight or wind, respectively. The duration of the thrips life cycle can range from 9 – 13 days from egg to adult depending on temperature (25°C – 30°C) and photoperiod (reviewed in Reitz, 2009). The short reproductive cycle and high fecundity of this species contributes to its success as an invasive species.

Fine-scale morphological documentation of egg structure and embryogenesis has been well documented for terebrantian thrips, including *F. occidentalis*. The eggs laid by *F. occidentalis* are pale white or light yellow in color, kidney-shaped, have a smooth outer layer (chorion) and possess an anterior operculum that is removed by the oviruptor, a serrated, knife-like structure, upon hatching of first-instar larvae. The width of the egg changes during development due to the accumulation of yolk over surrounding structures (Moritz, 1997). While eggs are naturally oviposited under the epidermis of plant tissues, egg-laying chambers that allow females to oviposit through artificial membranes sandwiched with sucrose solution or agar provide effective means for collecting large numbers of eggs for experimental purposes (Kumm, 2002; Moritz, 1997). The duration of *F. occidentalis* egg development is approximately four days at 23°C (Kumm, 2002), but can be reduced with increased temperature (Reitz, 2008). Shortly after oviposition, the nucleus of the zygote undergoes mitotic division, producing many daughter nuclei (syncytial blastoderm stage), and these resulting cleavage energids move to the egg periphery (plasmodial preblastoderm) (Kumm, 2002). Eggs reach the blastoderm stage a few hours after oviposition. Mitotic activity divides the blastoderm into the embryonic tissue and the extraembryonic cover, followed by anatrepsis when the head-end of the embryo is positioned at the posterior pole of the egg. The relatively rapid rate of thrips embryonic development and the requisite of precellular embryos for efficient editing of insects (Reid & O’Brochta, 2016) inform us that microinjection would need to occur within a few hours post oviposition for *F. occidentalis*.

Development of CRISPR-based genome editing in uncharted insect species typically aims to target a well-conserved gene known to have an overtly visible phenotype, such as those that influence eye color. All insect eyes studied to date possess ommochrome pigments (Grubbs et al., 2015), and disrupting ommochrome biosynthesis results in a change of pigmentation. This makes the genes involved in ommochrome biosynthesis ideal targets for testing Cas9 function. Two of these eye-color genes, *white* (*w*, encodes half of an ABC transporter) and *cinnabar* (*cn*, encodes kynurenine 3-monooxygenase), are the most popular targets in non-model insects. In *Drosophila*, the *w* gene product dimerizes with the *scarlet* or *brown* gene products to form complete ABC transporters that import ommochrome (brown) or pteridine (red) pigments into the cell, respectively (Dreesen et al., 1988; Sullivan et al., 1979; Sullivan & Sullivan, 1975; Tearle et al., 1989). Loss of *w* function usually leads to a white-eyed phenotype (Grubbs et al., 2015), in fact, *w* was among the first reports of Cas9-based gene knockout in a dipteran (Meccariello et al., 2017), two lepidopterans (Khan et al., 2017; Shirk et al., 2023), and several hemipterans (de Souza Pacheco et al., 2022; Klobasa et al., 2021; Reding & Pick, 2020; Xue et al., 2018). Ommochrome biosynthesis involves the catabolization of tryptophan into a series of intermediates that are processed into pigments. The kynurenine 3-monooxygenase protein encoded by *cn* is one of these essential enzymes involved in the tryptophan processing cascade (Linzen B, 1974). Recent reports of *cn* knockout in non-model insects includes western tarnished plant bug (Heu et al., 2022) and tomato leaf miner *Tuta absoluta* (Ji et al., 2022). Since loss of *w* can be lethal in some insect species (Khan et al., 2017; Klobasa et al., 2021; Perera et al., 2018), researchers working with non-model insects often aim to target more than one eye-color gene, as was the case of the first report of Cas9-mediated knockout of *w* and *cn* in the brown planthopper (Xue et al., 2018).

Here we report the first ‘proof-of-principle’ use of the CRISPR/Cas9 genome editing system in a thysanopteran. Suitable target genes, *w* and *cn*, were identified within the *F. occidentalis* genome and guide RNAs targeting each were generated. Precellular embryos were microinjected with Cas9 nuclease and guide RNAs targeting either *w* or *cn*. Injectees and their offspring were screened for changes in eye-color phenotype and putative mutants were confirmed by molecular and sequence analysis. This is the first step towards developing a new functional genomics tool for *F. occidentalis,* and for paving the way for sophisticated genetic methods that enable the use of gene-drive technologies for population suppression (local eradication) and/or population replacement (replace field populations with vector-incompetent thrips) as a means of mitigating virus transmission by this supervector.

## RESULTS AND DISCUSSION

### Identification of Frankliniella occidentalis white and cinnabar genes

Blast analysis using *N. lugens white* (ANO53447) as query against the *F. occidentalis* OGS revealed three high scoring matches. While the top match had the lowest e-value (0), the second match had a greater overall sequence identity (65% vs. 55%). The second match was shorter (i.e. partial sequence) but was referred to as *F. occidentalis white* (*Fo-w*, XM_052276619.1) in the NCBI database, while the top hit was named *white-like* (XM_052269207.1). The identities of these sequences were verified via BLASTX against the NCBI non-redundant (nr) protein database. The next highest matches shared the greatest sequence identities with the closely related eye-color genes, *scarlet* and *scarlet-like*. However, much further down the list (E-value = 2e-62) was another partial gene, which was referred to as *white-like* (XM_052277150.1). It was considerably shorter than the other sequences (∼1kb vs. >2kb) but appeared to possess the fragment that was missing in the previous *white* gene, and it shared a similar degree of sequence identity (62%) with the query. Together, the two partial sequences (XM_052276619.1 and XM_052277150.1) made up a nearly full-length *white* transcript.

To verify that the two partial transcripts came from the same locus and to determine gene structure, the mRNA sequences were used to search the *F. occidentalis* genome assembly. Although each sequence matched a distinct genomic scaffold, it was evident that they individually encoded separate parts of the same *white* gene. In fact, the genomic scaffolds (NW_026270734.1 and NW_026272940.1) revealed the four nucleotides required to assemble the two partial transcripts together (data not shown). In addition, comparison of the full-length transcript to the super-scaffold revealed that *Fo-w* consists of 11 coding exons.

Similarly, using *N. lugens cinnabar* (ALQ52680.1) as query against the *F. occidentalis* OGS identified a partial mRNA (XM_052275949.1), which was labeled as kynurenine 3-monooxygenase. The identity of the partial mRNA was verified via BLASTX analysis in the NCBI nr protein database but the 5’-most coding sequence was found to be missing. To discover the 5’-most end of *F. occidentalis cinnabar* (*Fo-cn*) gene, the partial mRNA sequence was employed to identify the corresponding genomic scaffold. Although sequence analysis revealed a single genomic scaffold (NW_026268346.1), it failed to provide new insights into the missing 5’ end of the gene due to the relatively small size of the scaffold in question (<12kb). Therefore, we used the deduced amino acid sequence of the full-length *cinnabar* transcript (XM_034398812.1) from a closely related thrips species, *Thrips palmi*, to determine if the genome assembly possessed a scaffold that could potentially encode the 5’ region of *Fo-cn*. This analysis led to the identification of another *F. occidentalis* genomic scaffold (NW_026267160.1) which appears to encode the 5’ end of *Fo-cn* gene. A total of nine exons were identified by comparing the full-length *Fo-cn* transcript to the super-scaffold.

### CRISPR/Cas9-mediated knockout of *F. occidentalis white* and *cinnabar* genes

Due to the essential role of *white* and *cinnabar* in the biosynthesis and transport of ommochrome and pteridine pigments, as well as the well-documented correlation between loss-of-function mutations in the *white* and *cinnabar* genes and the complete loss or alteration of eye pigmentation in many insect species (Ewart & Howells, 1998; Mackenzie et al., 1999, 2000; WD et al., 1996), we sought to test CRISPR-mediated genome editing in thrips by targeting *Fo-w* and *Fo-cn*. Among the 104 embryos injected with single-guide (sg) RNAs targeting *Fo-w*, 25 hatched, with 17 of the larvae displaying varying degrees of eye-color knockout (**Table 1**, **Figure 1B-C**). Since high rates of somatic mutation in G_0_ individuals is correlated with higher rates of gene edited germline cells (Han et al., 2023), we focused on the two female and two male G_0_ adults that displayed a red-eye phenotype. Progeny from self-crossing these four individuals were used to establish a homozygous *Fo-w* knockout line (**Figure S1**).

**Figure 1.**
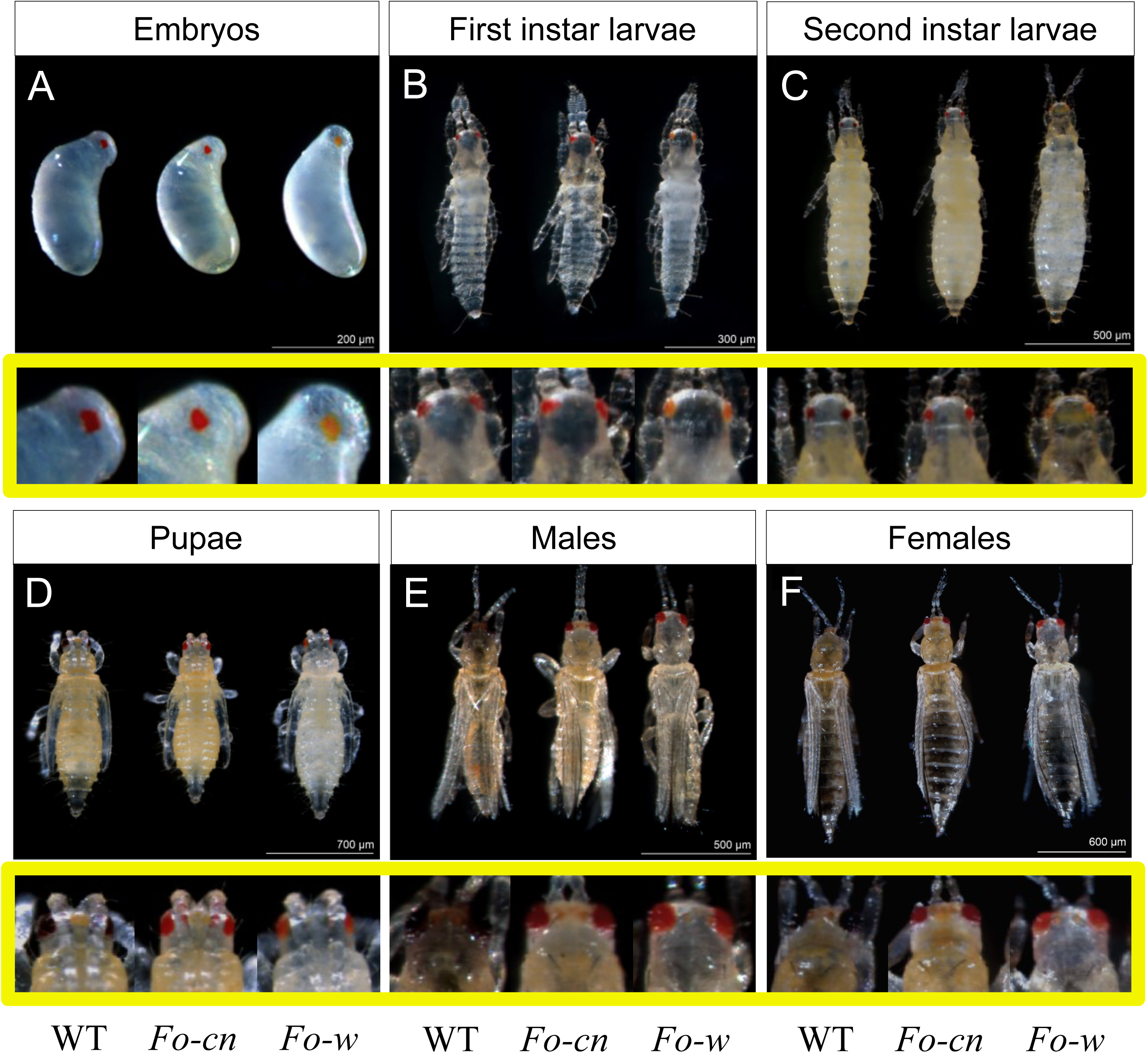
Eye-color phenotypes of *Fo-w* and *Fo-cn* knockout lines across the different developmental stages of *Frankliniella occidentalis.* Phenotypic changes in thrips eye color were photographed in embryos (A), first-instar larvae (B), second-instar larvae (C), pupae (D), adult males (E), and adult females (F) of knockout lines compared to wild type (WT). Panels within the yellow squares are magnified photos for a better visualization of thrips eye pigmentation. Photographic specimens were not age synchronized and were selected based on eye-color phenotype. Other phenotypic differences between specimens, such as body color or size, are not representative of their respective genotype.

**Table 1:** Numbers of surviving embryos and eye-color gene (*Fo-w*, *Fo-cn*) knock-out phenotypes in *Frankliniella occidentalis* after embryo microinjection.

| Metric | White | Cinnabar |
| --- | --- | --- |
| Total number of injected embryos | 104 | 53 |
| Number of hatched larvae from injected embryos | 25 | 16 |
| Hatching rate of embryos | 24% | 30% |
| *Number of larvae with distinguishable eye-color knockout | 17 | 0 |
| Number of knockout adult females | 2 | 2 |
| Number of knockout adult males | 2 | 1 |
| Knockout rate of adults | 3.9% | 3.8% |

\*The eye pigmentation of cinnabar gene knockout larvae in the G<sub>0</sub> generation showed no discernible difference from the wild type. Notably, a distinctive alteration in eye color was observed specifically in G<sub>0</sub> adults.

Of 53 embryos injected with sgRNAs targeting *Fo-cn*, 16 hatched; however, none of the larvae displayed a discernible phenotype (**Table 1**). Interestingly, upon reaching adulthood, three (two females and one male) showed a knockout phenotype. However, self-crossing the three phenotypic thrips failed to produce progeny having the expected homozygous *Fo-cn* knockout phenotype (**Figure S2**). The remaining 13 non-phenotypic G_0_ adults were also allowed to self-cross and in the subsequent generation, a total of 28 G_1_ individuals displaying mutant eye-color phenotypes were identified. These G_1_ individuals were used as founders for establishing a stable, homozygous *Fo-cn* knockout line (**Figure S1**).

Due to the importance of eye-color genes as markers for large-scale mutagenic screens, a technique known as co-CRISPR (Kane et al., 2017), we have given much thought to the outcomes of targeting *Fo-w* vs. *Fo-cn*. Specifically, somatic mosaic knockout phenotypes for *Fo-w* proved to be an excellent indicator for finding mutant G_1_ offspring. On the other hand, using somatic mosaic knockout phenotypes for *Fo-cn* as an indicator failed. The difference is likely due to the fact that *Fo-cn* encodes a cell non-autonomous protein which can move from cell to cell. This is not true for *Fo-w*. Therefore, in the case of *Fo-cn*, G_0_ individuals could have a large number of mutant cells but nonetheless still have enough wild-type protein to make wild-type eye pigments. We conclude that targeting a gene that encodes a cell-autonomous protein like *Fo-w* will provide a better indicator of potential germline change than those that encode cell non-autonomous proteins. While somatic knockout phenotypes in injectees are only an indicator that germline cells might also carry the desired mutation, decreasing the number of G_0_ individuals that have to be followed, thus G_1_ offspring that have to be screened, can dramatically increase the efficiency of large-scale screens aimed at finding knockout alleles for genes that lack a known phenotype, or screens aimed at finding rare events such as homology directed repair mediated knock in.

### Phenotypes of *Fo-w* and *Fo-cn* knockout thrips lines

Thrips eye-color changes were documented throughout the different developmental stages and among both sexes of *Fo-w* and *Fo-cn* homozygous knockout lines, including embryo, first instar larva, second instar larva, pupa, adult female, and adult male (**Figure 1**). However, the *Fo-w* knockout phenotype was not that of complete loss of pigmentation, rather larvae possessed ivory-colored eyes in contrast to the dark red color observed in wild type (WT) (**Figure 1B-C**). Moreover, as the individuals aged, eye pigment changed from ivory to bright red (**Figure 1**). Considering that male thrips are haploid, any male possessing a mutated *white* allele was expected to have white eyes. However, the observed ivory eyes in *Fo-w* mutant males suggests that the thrips eye has some degree of pigmentation despite loss of *white* gene function. Even more unexpected was the fact that their eye color changed to a bright red as they aged. The mutant phenotype was still clearly distinct from that of WT adults enabling selection of mutant males to cross with red-eyed females, but the question remains as to why their eye color changes from ivory to red as they age.

During colony maintenance we noted that body pigmentation appeared to be lighter in color for individuals from the *Fo-w* knockout lines than that observed for wildtype thrips under the same environmental, rearing conditions. Sampling, visual assessment and statistical analysis of body pigmentation (unpigmented (white), partially pigmented, amber pigmented) of individuals in groups of 100 L2 thrips per genotype (*Fo-w* knockout and wildtype) revealed an overall significant effect of genotype on body-color phenotype (Fisher Exact test, two-sided p-value < 0.0001). For the *Fo-w* knockout sampling set, 63.3% exhibited unpigmented bodies and 3% amber-body coloration, while in the wildtype set, only 7% exhibited unpigmented bodies and 49% presented amber color (**Figure 2C**). Fisher Exact tests on contingency (frequency) tables for all possible pairwise phenotype comparisons (three) revealed significant differences due to genotype (two-sided p-value < .0001), including the patchy coloration compared to the amber (complete) coloration. This alteration in thrips’ body pigmentation is likely an unintended consequence of disrupting ommochrome or pteridine pigment synthesis. These pigments can not only affect eye color but also body color, as observed in other arthropods (Kômoto et al., 2009; Liu et al., 2021; Reding & Pick, 2020). While not all arthropods rely on these pigments for body coloration, this initial data suggests that *F. occidentalis* likely relies on pteridines for body pigmentation (i.e. loss of ommochrome pigment pathway lacks visible difference in body color). It is important to note, however, that *F. occidentalis* can exhibit various color morphs that are influenced by environmental conditions (O’Donnell, 2006). Despite this, all *F. occidentalis* colonies in this study were reared under controlled conditions that would not typically yield such body color differences. Future experiments aiming to comprehensively characterize the effects of eye color mutations on body pigmentation would involve further documentation of the frequency of lightly pigmented thrips across various life stages. Moreover, quantitative studies using high-performance liquid chromatography (HPLC) can be conducted to measure differences in the abundance of pteridine pigments and their precursors between wild type and eye-color mutant thrips.

**Figure 2.**
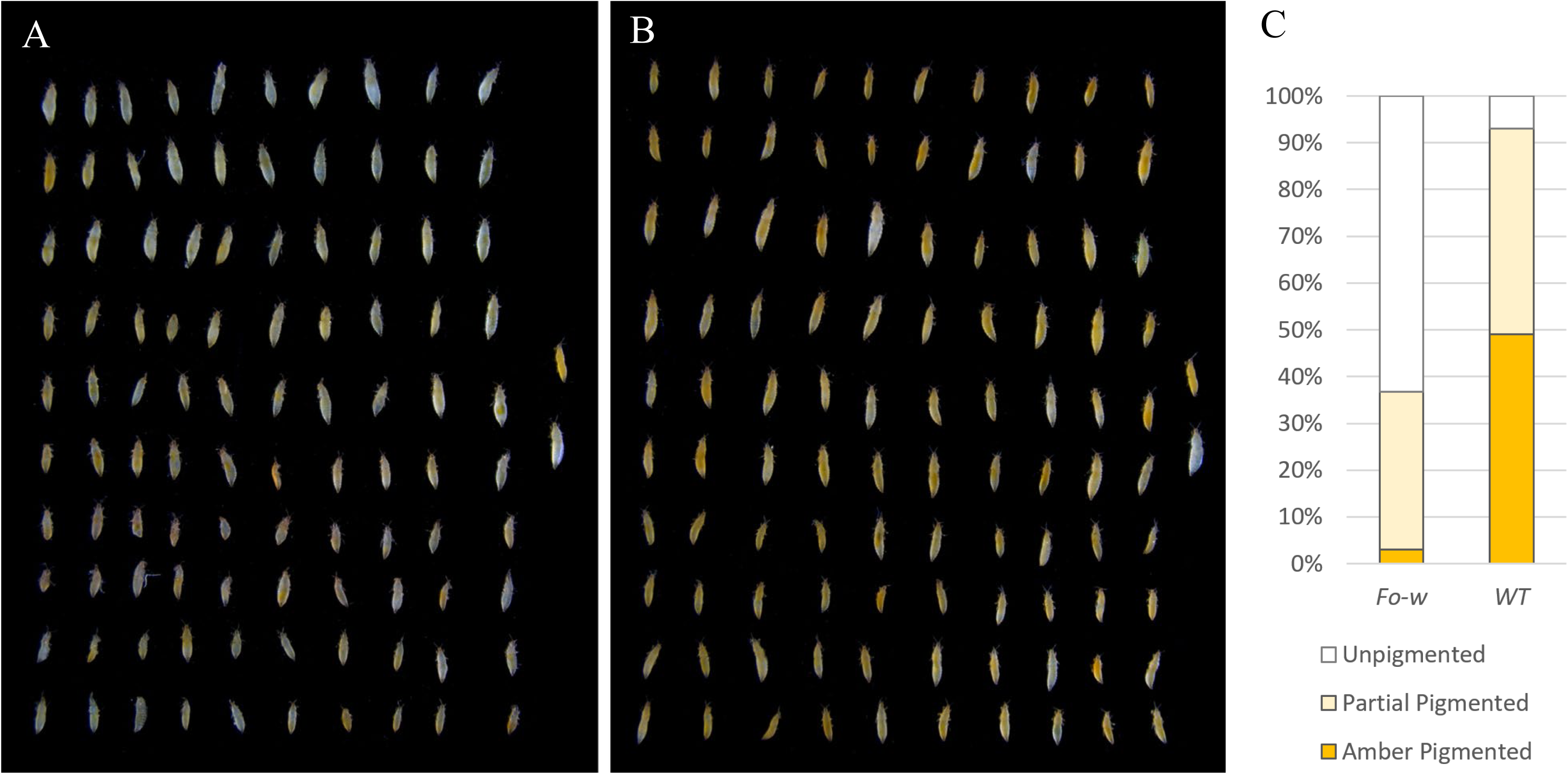
Visual assessment of *Frankliniella occidentalis* second-instar larval body-color phenotypes of (A) *Fo-w* knockout and (B) wild-type (WT) specimens. Thrips larval specimens were randomly collected from both the *Fo-w* knockout line and WT to create separate 10 x 10 grids. The two specimens depicted on the right side of (A) and (B) are identical, serving as a reference for evaluating the color-morph phenotype (non-pigmented, partial-pigmented, or pigmented) of the remaining specimens. Individuals exhibiting a lack of pigmentation similar to the bottom reference specimen were classified as non-pigmented; those that had inconsistent splotches of amber pigmentation were classified as patchy-pigmented; and those that had consistent amber pigmentation similar to the top reference specimen were classified as pigmented. (C) Percentages of each color morph between the *Fo-w* and WT knockout mutant specimens (Fisher Exact Test of Independence, two-sided p-value < 0.0001).

As noted above, *F. occidentalis* has clear duplications of the ABC transporters known to be involved in eye pigmentation. If the duplicated genes serve the same function, and possess the same expression patterns, a mutant phenotype would only be expected to be seen when both genes are disrupted. However, since a mutant phenotype (ivory-colored eyes) was observed in late-stage embryos (**Figure 1A**) and during both larval instars (**Figure 1B-C**), the most likely explanation is that duplication of *white*, giving raise to *white-like*, generated a version of *white* that serves the original function, but its expression is delayed until the pre-pupal or pupal stage of development. If this stands to be the case, both *white* and *white-like* would need to be knocked out to establish a white– or ivory-eyed line. Importantly, since the deletion in the *Fo-w* knockout line results in complete loss of the region that encodes the Walker B motif (CDEPT), it seems highly unlikely that the mutant allele has partial functionality. Future studies to test the hypothesis that microinjection of guides specific for *white-like* into embryos derived from the *Fo-w* knockout line would produce a true white-eyed line are warranted.

Unlike the *Fo-w* knockout line, which displays an age-dependent knockout phenotype, the *Fo-cn* knockout line exhibited the same phenotype throughout all life stages, that of bright red eyes (**Figure 1**). Similar knockout phenotypes have been reported for *cn* in other insect systems (Heu et al., 2022; Sullivan et al., 1979; Vargas-Lowman et al., 2019; Xue et al., 2018). For example, while the brown planthopper eye usually appears brown, mutagenesis of the *Nl*-*cinnabar* gene led to a striking transformation to bright red (Xue et al., 2018).

### Molecular and sequence confirmation of *Fo-w* and *Fo-cn* knockout in thrips

Cas9-mediated knockout of *Fo-w* and *Fo-cn* was confirmed through PCR and agarose gel electrophoresis, which demonstrated that the amplification products derived from DNA isolated from mutants obtained from the respective knockout lines were significantly smaller compared to that from WT thrips. Specifically, amplification products generated from the *Fo-w* knockout line were consistent with a deletion of ∼732 bp, the size of the genomic region between the *Fo-w* sgRNAs located in exon #2 and exon #4 (**Figure 3A**). Similarly, the amplification products derived from DNA isolated from the *Fo-cn* knockout line suggests a deletion of ∼633 bp, the size of the genomic region between the *Fo-cn* sgRNAs located in exons #1 and #4 of the *cinnabar* gene (**Figure 3B**). This indicates that the targeted regions of these genes were successfully removed from the thrips genome.

**Figure 3.**
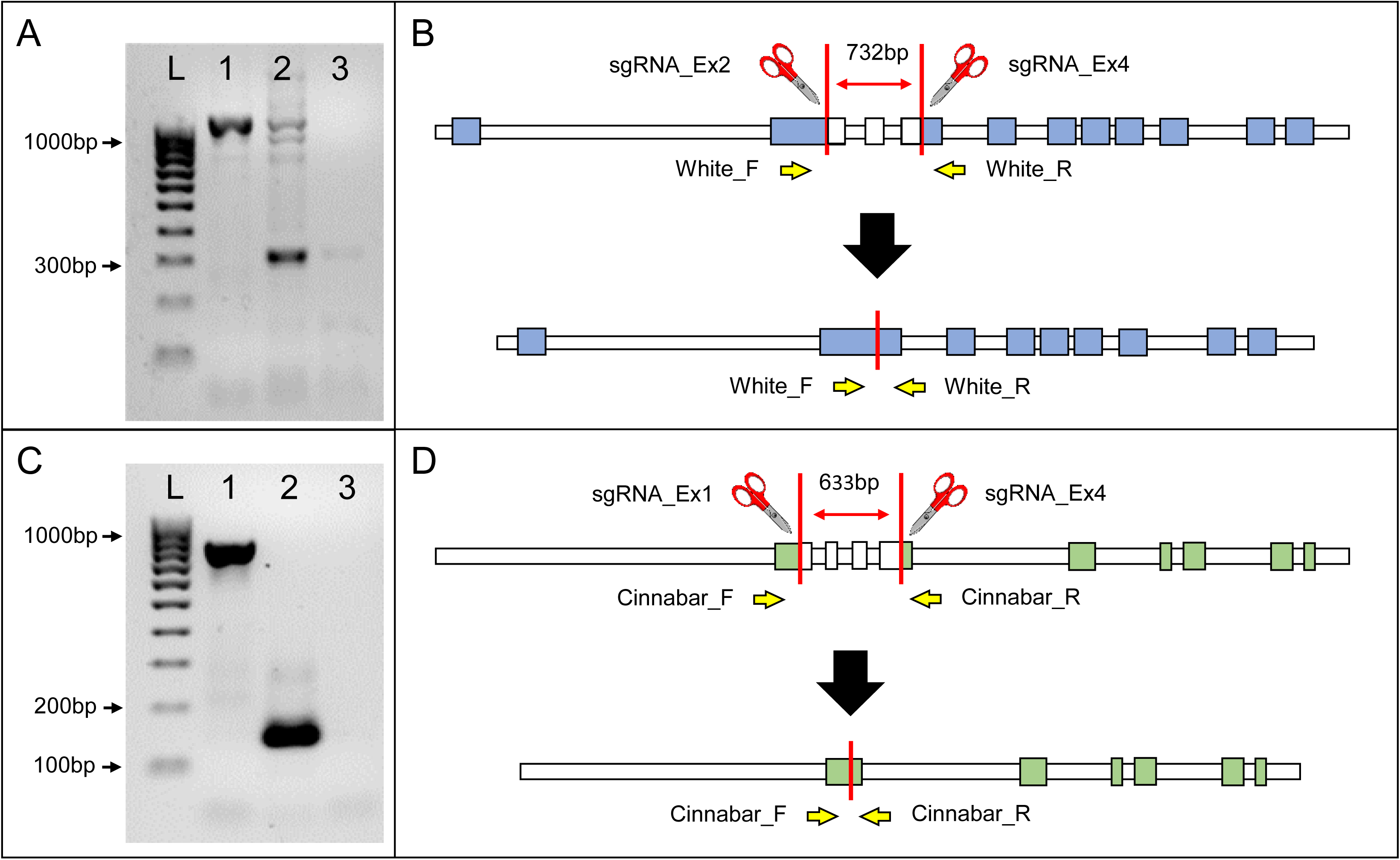
Knockout of *Fo-w* and *Fo-cn* eye-color genes of *Frankliniella occidentalis*. Agarose gel resolutions of CRISPR-mediated deletions of target nucleotide sequences in (A) *Fo-w* and (B) *Fo-cn* knockout G_1_ adults, respectively. Comparison of Lane 1 and 2 demonstrates the removal of 732bp and 633bp of wild-type (WT) sequence within the *white* (*w*) and *cinnabar* (*cn*) genes, respectively. L = ladder; 1 = WT template DNA; 2 = *Fo-w*/*cn* G_1_ DNA; 3 = no DNA template control. (B) and (D) Scale representations of WT *Fo-w* and *Fo-cn* genes and their corresponding CRISPR-mediated deletions. Exons are shown in blue or green boxes. The scissors and white boxes with diagonal stripes indicate cut sites and regions within the exons. PCR primer sites are depicted by yellow arrows.

Moreover, we validated proper Cas9-mediated editing by sequencing the amplification products. As expected, pairwise alignments of DNA sequence derived from the *Fo-w* knockout line with that from WT thrips revealed a sizable deletion, consistent with Cas9 cleavage at the intended sites. For example, the pairwise alignment shown in **Figure 4** indicates that Cas9 induced double-strand breaks at positions 194 bp and 926 bp, resulting in a loss of 732 bases. Similarly, the pairwise alignment of sequence data from the *Fo-cn* knockout line (**Figure 5**) and WT also demonstrates Cas9-induced deletion of the sequence located between the sgRNA cut sites (positions 94 bp and 727 bp). These findings affirm the feasibility of sequence-specific gene knockout in thrips using CRISPR-mediated genome editing.

**Figure 4.**
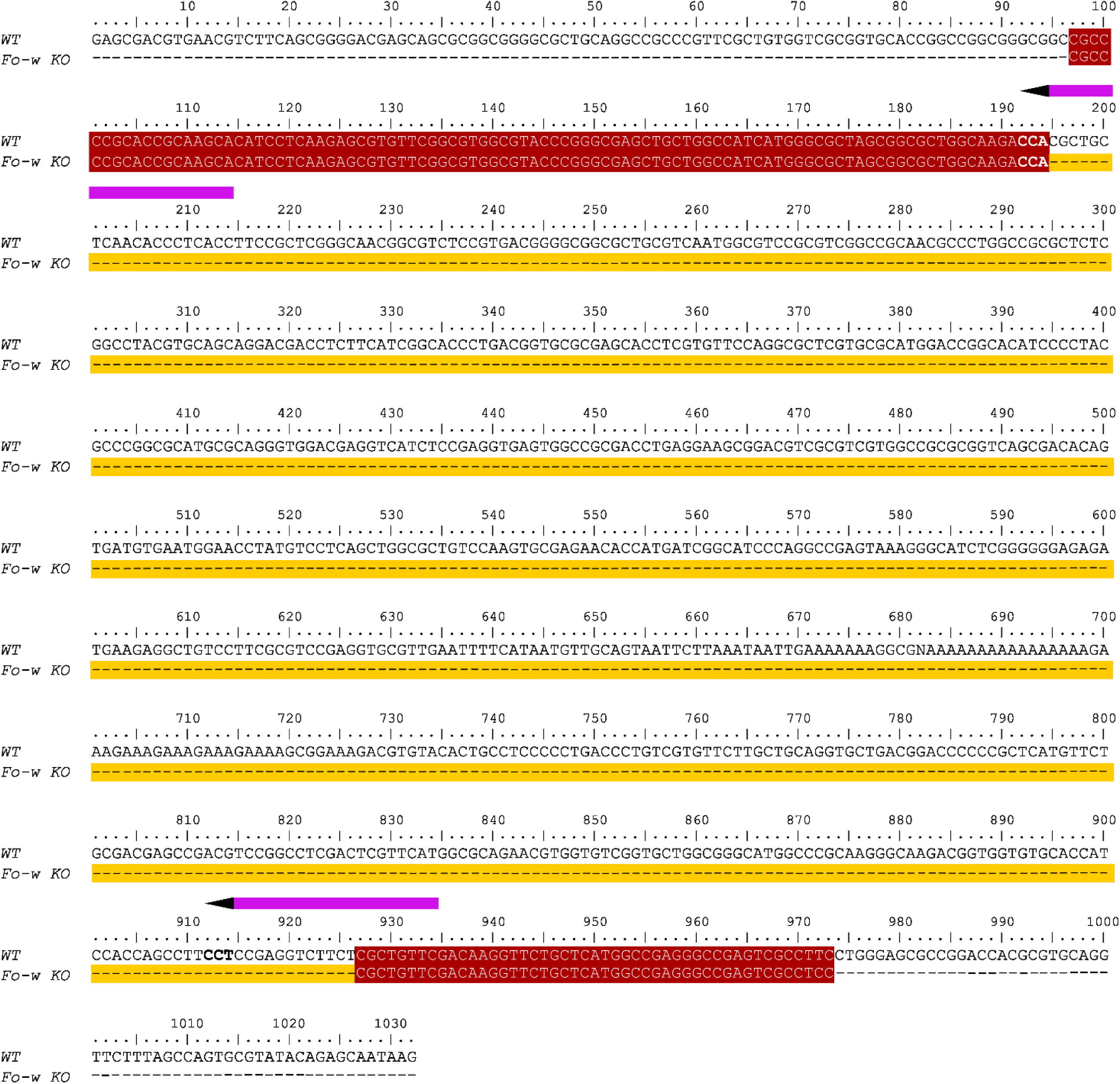
Pairwise sequence alignment of *Fo-w* gene between wild type (WT) and *Fo-w* knockout (KO) thrips of *Frankliniella occidentalis*. A 732bp nucleotide sequence deletion was confirmed in the genome of *Fo-w* KO thrips compared to WT (highlighted in yellow). Aligned sequences are highlighted in red color. The pink arrow bars above the alignment indicate the two single guide RNAs targeting the *white* gene. The direction and head of arrows represents the RNA strand from 5’ to 3’ and the protospacer adjacent motif (PAM).

**Figure 5.**
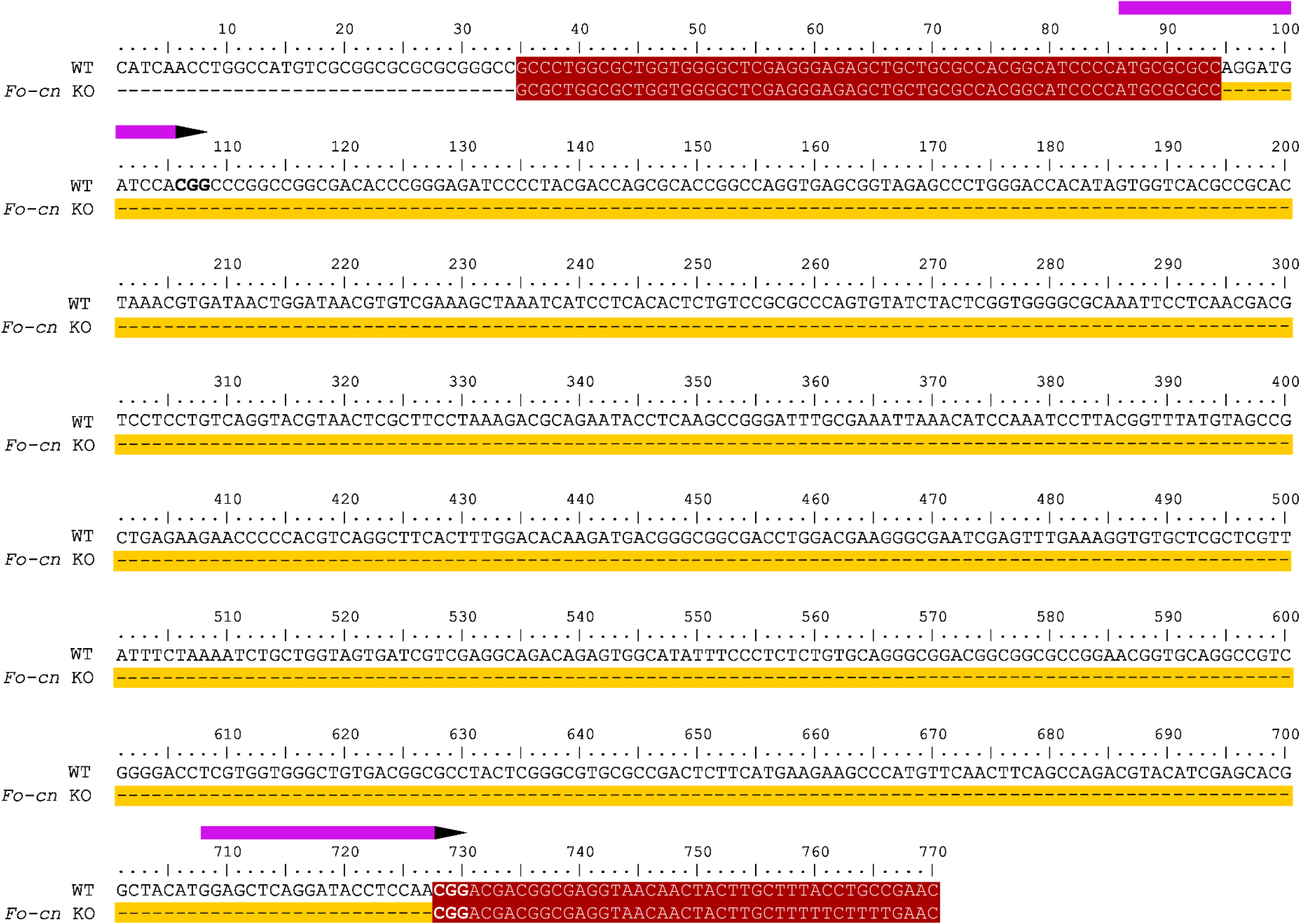
Pairwise sequence alignment of *Fo-cn* gene between wild type (WT) and *Fo-cn* knockout (KO) thrips of *Frankliniella occidentalis*. A 633bp nucleotide sequence deletion was confirmed in the genome of *Fo-cn* KO thrips compared to WT (highlighted in yellow). Aligned sequences are highlighted in red color. The pink arrow bars above the alignment indicate the two single guide RNAs targeting the *cinnabar* gene. The direction and head of arrows represents the RNA strand from 5’ to 3’ and the protospacer adjacent motif (PAM).

Overall, our study describes a simple and effective method for generating heritable mutations in the *F. occidentalis* germline using the CRISPR/Cas9 system. It not only offers an alternative approach to investigate the function of thrips genes, but also complements the existing RNA interference techniques developed for thrips (Maurastoni et al., 2023). Notably, given the economic significance of thrips as a pest, the successful implementation of CRISPR/Cas9-mediated genetic manipulation holds promise for controlling thrips populations in real-world field applications using sophisticated genetic techniques aimed at population replacement, or population suppression (Esvelt et al., 2014; Kandul et al., 2019). Of these, gene drive is particularly useful for replacing populations of virus-competent insects with those that can no longer vector viruses. Most examples involve generation of gene drive mosquitoes aimed at reducing transmission of malaria (reviewed in James, 2005; Nawy, 2016). To date, genetic control of agricultural pests is mainly through the use of sterile insect technique (SIT; Knipling, 1959), with eradication of the New World screwworm, *Cochliomyia hominivorax*, from the continental US being the biggest success story (Wyss, 2000). However, standard SIT employs isotopes to sterilize insects which can reduce their fitness. Therefore, researchers turned to CRISPR/Cas9 to improve the system. The CRISPR-based system is known as precision-guided SIT (pgSIT) and sterilizes insects by modifying genes responsible for fertility (Kandul et al., 2019) in the model system, *Drosophila melanogaster* (Kandul et al., 2019, 2021), as well as non-model species outside ofdrosophilids (Akbari et al., 2023). Demonstration of CRISPR/Cas9 efficacy in *F. occidentalis* in the present proof-of-principle study enables the development and enhancement of novel biotech-based methodologies for controlling thrips vectors and the viruses they transmit to plants (Maurastoni et al., 2023).

## EXPERIMENTAL PROCEDURES

### Identification of two *Frankliniella occidentalis* eye-color genes and CRISPR guide design

The official gene set (Focc_3.1) of the *F. occidentalis* genome (NCBI RefSeq assembly: GCF_000697945.3) was downloaded and queried locally (BlastStation, TM Software, Arcadia, CA) for putative orthologs of *w* and *cn* eye-color genes with *Nilaparvata lugens* (brown planthopper) white (ANO53447) and cinnabar (ALQ52680) amino acid sequences using TBLASTN analysis (Altschul et al., 1997). The resulting top-scoring matches for each gene were analyzed using NCBI BLASTX to confirm their annotations, and the corresponding genomic scaffolds were located within the thrips genome (PRJNA203209) using BLASTN.

To enable robust single guide RNA (sgRNA) design for *w* and *cn* genes identified for *F. occidentalis* (designated herein as *Fo-w* and *Fo-cn*), two different programs were used – IDT (https://www.idtdna.com/site/order/designtool/index/CRISPR_CUSTOM) and CRISPOR (http://crispor.tefor.net/crispor.py) – and candidate sgRNA sequences found to be in common between both program outputs were selected for further analysis. To improve the specificity and efficiency of gene knockout, two sgRNAs were designed for targeting two exons per eye-color gene: exon 2 and 4 of *Fo-w* and exon 1 and 4 of *Fo-cn*. The specificity of each sgRNA was evaluated via BLASTN analysis against the i5K *F. occidentalis* genome assembly (Rotenberg, Gibbs, et al., 2019; Rotenberg, Robertson, et al., 2019) and the NCBI RefSeq gene set database (GCF_000697945.3). To minimize off-target cutting, sgRNAs matching other non-target sequences were required to have at least five nucleotides of mismatch at the 3’ end to remain on the list of candidates. All sgRNAs were synthesized by Synthego (Redwood City, CA), with modification to add 2’-O-Methyl at the three first and last bases, and 3’ phosphorothioate bonds between the first three and last two bases. The final sgRNA sequences used in this study are listed in **Table S1**.

### Microinjection of *F. occidentalis* embryos

The *F. occidentalis* colony used in this work was originally collected from the island of Oahu, HI, USA and maintained on green bean pods (*Phaseolus vulgaris*) at 25°C, 16L:8D photoperiod as described previously (Ullman et al., 1992). Precellular embryos were collected from approximately 500 age-synchronized adult thrips (2 – 12 days after eclosion) that were allowed to oviposit for three hours on a stretched Parafilm membrane into a 10-cm Petri dish containing 3% sucrose solution. The embryos were collected using a coffee paper filter and rinsing with distilled water. Wet filter paper carrying the embryos was gently transferred into a 15 cm Petri dish and covered with a lid to maintain high humidity. Prior to microinjection, 10 – 30 embryos were gently transferred onto a strip of double-sided tape on a microscope slide using a fine paint brush under a dissecting microscope. The posterior end of each embryo was positioned towards the microinjection needle.

All sgRNAs targeting *Fo-w* and *Fo-cn* were diluted into a working solution of 400 ng/μl with a nuclease-free 1X Tris-EDTA buffer. The injection mixture was prepared by complexing 0.5 μl of Invitrogen TrueCut Cas9 protein (714 ng/ul) (Thermo Fisher Scientific Inc., Waltham, MA) with 1.5 μl of each of the two sgRNAs. The mixture was gently mixed by tapping the bottom of the tube and centrifuged for a few seconds. For proper complexing, the mixture was incubated at room temperature for 15 minutes and then placed on ice before use. The mixture (1 μl) was pipetted into a tapered, quartz glass needle (od=1.0; id=0.5) using a microloader tip (Eppendorf, Hamburg, Germany) and inserted into a needle holder of an in-house microinjection system supported by an Eppendorf Femtojet 5247 programmable microinjector pump (Eppendorf, Hamburg, Germany). Each embryo was injected with the sgRNAs/Cas9 mixture through its posterior pole and each cohort of injected embryos was immediately moved into a Petri dish containing a thick layer of agarose gel in a humidified incubation chamber and reared at 25°C. Approximately 48 hours post injection, the embryos were carefully transferred from the sticky tape to a Petri dish containing a moistened filter paper. After hatching, individuals were transferred to rearing containers filled with green bean pods and monitored daily for eye-color phenotypes. A detailed microinjection protocol is described in **Supplementary File 1**.

### Establishment of homozygous *Fo-w* and *Fo-cn* knockout lines

Less than 24h post eclosion, G_0_ adults were screened for phenotypic changes in eye color under a dissecting microscope. Individuals having *Fo-w* or *Fo-cn* mutant phenotypes were transferred into new rearing containers filled with green bean pods and allowed to mate. The resultant G_1_ progeny were examined across the different developmental stages for eye-color phenotypes. Homozygous lines were established by backcrossing mutant G_1_ males with G_0_ females. Given the haplodiploidy characteristic of *F. occidentalis* (haploid males arise from unfertilized eggs, while diploid females originate from fertilized eggs) (Moritz, 1997) along with the recessive nature of the eye-color mutations, this crossing scheme provided the most straightforward mechanism for establishing homozygous lines. Specifically, screening for knockout events in haploid G_1_ males negated the possibility of a wild-type allele masking a mutated allele as is the case with diploid females in the absence of biallelic knockout. Similarly, backcrossing mutant G_1_ males to G_0_ females also negated the need for biallelic knockout since G_1_ females would inherit a mutant allele from their fathers. Thus, phenotypic G_1_ males were allowed to cross with non-phenotypic G_0_ females. Their progenies were screened for changes in eye color and those showing a change in phenotype were transferred to a new container to establish a homozygous line.

### Body pigmentation in *Fo-w* knock-out lines

One unexpected phenotypic outcome of the *Fo-w* gene knockout was a notable difference in body pigmentation. While maintaining the *Fo-w* knockout colony, we observed that all life stages beyond the first instar exhibited a lighter or patchy cuticle pigmentation in contrast to the amber cuticle typically observed in our wildtype colonies. To investigate whether this observed variation in body color could be attributed to genotype (knockout vs wildtype), we conducted a power analysis using R Studio (effect size = 0.3, alpha = 0.05, power = 0.80). This analysis aimed to calculate the appropriate sample size, determining the number of individuals to be sampled from each colony for a subsequent test of independence between phenotype (non-pigmented, semi-pigmented, or pigmented) and genotype (knockout vs wildtype). For each genotype, we sacrificed one colony cup enriched with second-instar larvae. Once immobilized, we arbitrarily sampled 100 second-instar larvae (L2) under magnification, adjusting the brightness settings on the microscope stage to make cuticle color indistinguishable. The specimens were then arranged into two 10 x 10 grids (**Figure 2A-B**) for visual assessment and enumeration of individuals classified into each color type. Fisher Exact Tests of independence on a two (genotype) by three (phenotype) contingency table was performed using JMP Pro 15 (SAS Inc), followed by pairwise comparisons between phenotypes using reduced contingency tables (2 by 2).

### Molecular and sequence analyses of gene knockout in thrips

Genomic DNA (gDNA) was isolated from two G_1_ adults per mutant line using Zymo Quick-DNA MicroPrep kit (Zymo Research, Irvine, CA), and the yield was quantified using a NanoDrop ND-1000 spectrophotometer (Thermo Fisher Scientific Inc., Waltham, MA). Putative genomic deletions in the *Fo-w* and *Fo-cn* knockout lines were evaluated by amplifying 12 ng of gDNA in 50 μl PCR reactions using a C1000 Touch Thermal Cycler (BioRad, Berkeley, CA). All reactions were carried out to the manufacturer’s recommendations using Ex Taq polymerase (Takara Bio, Shiga, Japan) and primer pairs (**Table S2**) designed to span the target region (Primer3Plus; Untergasser et al., 2007). Amplification of the *Fo-cn* deletion required an additional round of “nested” PCR using 1.5 μl of PCR product as a template. PCR amplicons were directly sequenced by Eton Bioscience (Durham, NC). A pairwise sequence alignment was performed to compare the genomic regions of *Fo-w* and *Fo-cn* obtained from the knockout lines to that of wildtype. The sequenced PCR amplicons of knockout thrips were aligned against the genomic sequence of wildtype using MUSCLE in MEGA11 software (Tamura et al., 2021).

## SUPPLEMENTARY MATERIALS

**<u>Supplementary File 1</u>**. Detailed microinjection protocol for *Frankliniella occidentalis* embryos (PDF)

**<u>Figure S1</u>**. Crossing scheme for establishing *white* and *cinnabar* mutant lines following G0 embryo injection. (PDF)

**<u>Figure S2</u>**. Genetic explanation for the failure and success of establishing *cinnabar* mutant line. A) Blue squares show the potential events occurred to produce non-phenotypic heterozygous progenies. (PDF)

**<u>Table S1</u>.** Single guide RNA (sgRNA) designed for CRISPR-mediated knockouts of *white* and *cinnabar* genes in *Frankliniella occidentalis.* (DOC)

**<u>Table S2.</u>** Primer pairs designed to detect the genomic sequence deletions in *white* and *cinnabar* knockout lines of *Frankliniella occidentalis.* (DOC)

## Author Contributions

Conceptualization, A.E.W., D.R and M.L; methodology, M.L., W.K, J.H., and L.dO.; formal analysis, J.H. W.K. M.L., L.dO.; investigation, J.H., W.K., L.dO.; resources, D.R. and M.L.; data curation, J.H. and M.L.; writing-original draft and editing, J.H., D.R., M.L., L.dO., W.K. and A.E.W.; visualization, J.H., W.K., and L.dO.; supervision, D.R. and M.L.; project administration, D.R. and M.L.; funding acquisition, M.L., D.R. and A.E.W. All authors have read and agreed to the published version of the manuscript.

## Data Availability

All sequence and database files used in the present study provided and/or cited in the text; any biological data not provided is available upon reasonable request.

## Supporting information

Supplementary File 1 Microinj Protocol

## Acknowledgements

We thank George Kennedy for sharing his procedure on thrips egg collection. We also thank the many undergraduate lab assistants of the Plant Virus Vector Interaction Lab at NCSU who contributed time and effort to maintaining a healthy and productive *F. occidentalis* colony over the course of this study.

