## Supplementary Figures S1 and S2 for "CRISPR/Cas9-mediated genome editing of *Frankliniella occidentalis,* the western flower thrips, via embryonic microinjection"

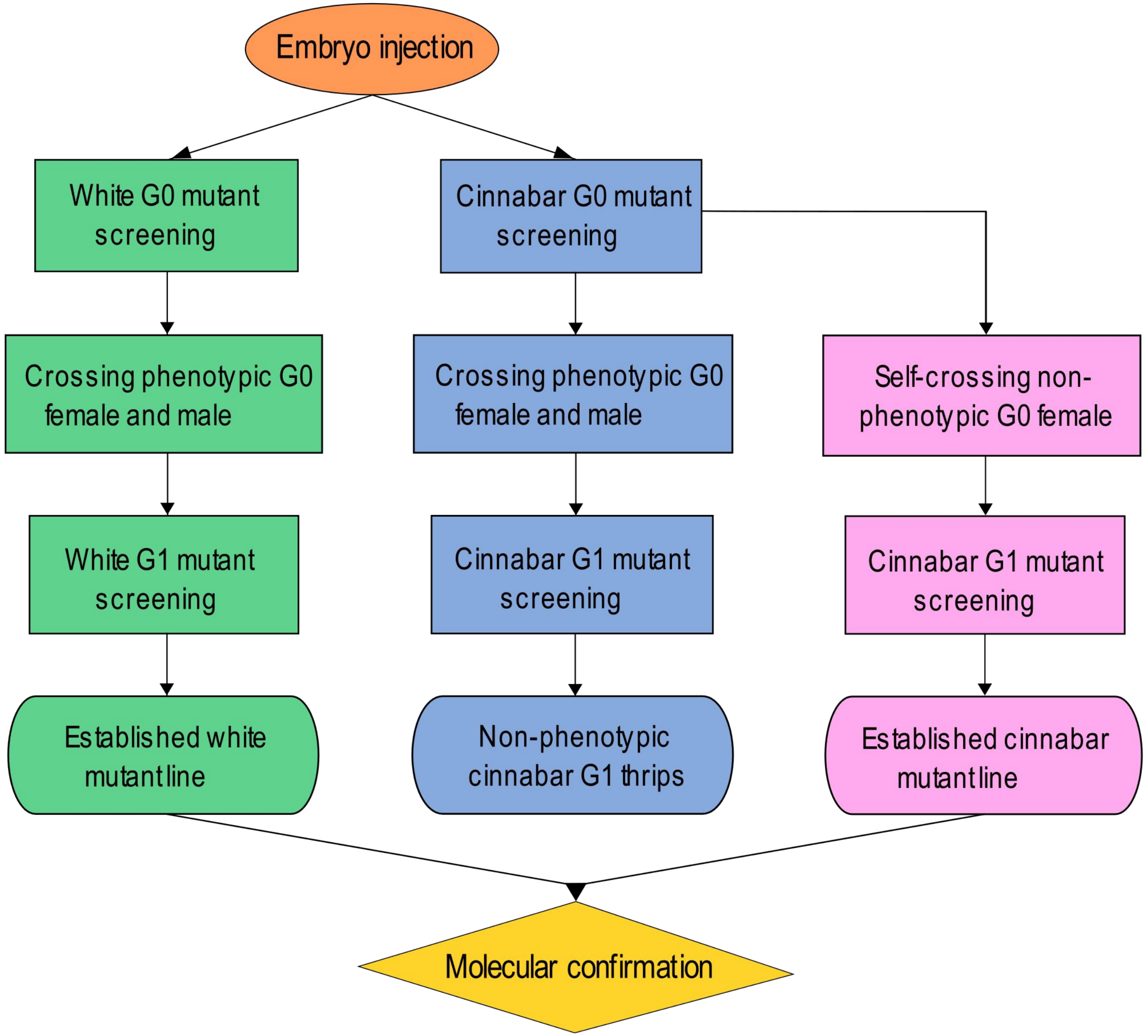


**Figure S1.** Crossing scheme for establishing *white* and *cinnabar* mutant lines following G0 embryo injection. Screening steps were represented by green squares for white mutant line and blue and pink squares for cinnabar mutant line. Blue and pink squares indicate the first and second attempts to establish a homozygous phenotypic cinnabar mutant line.

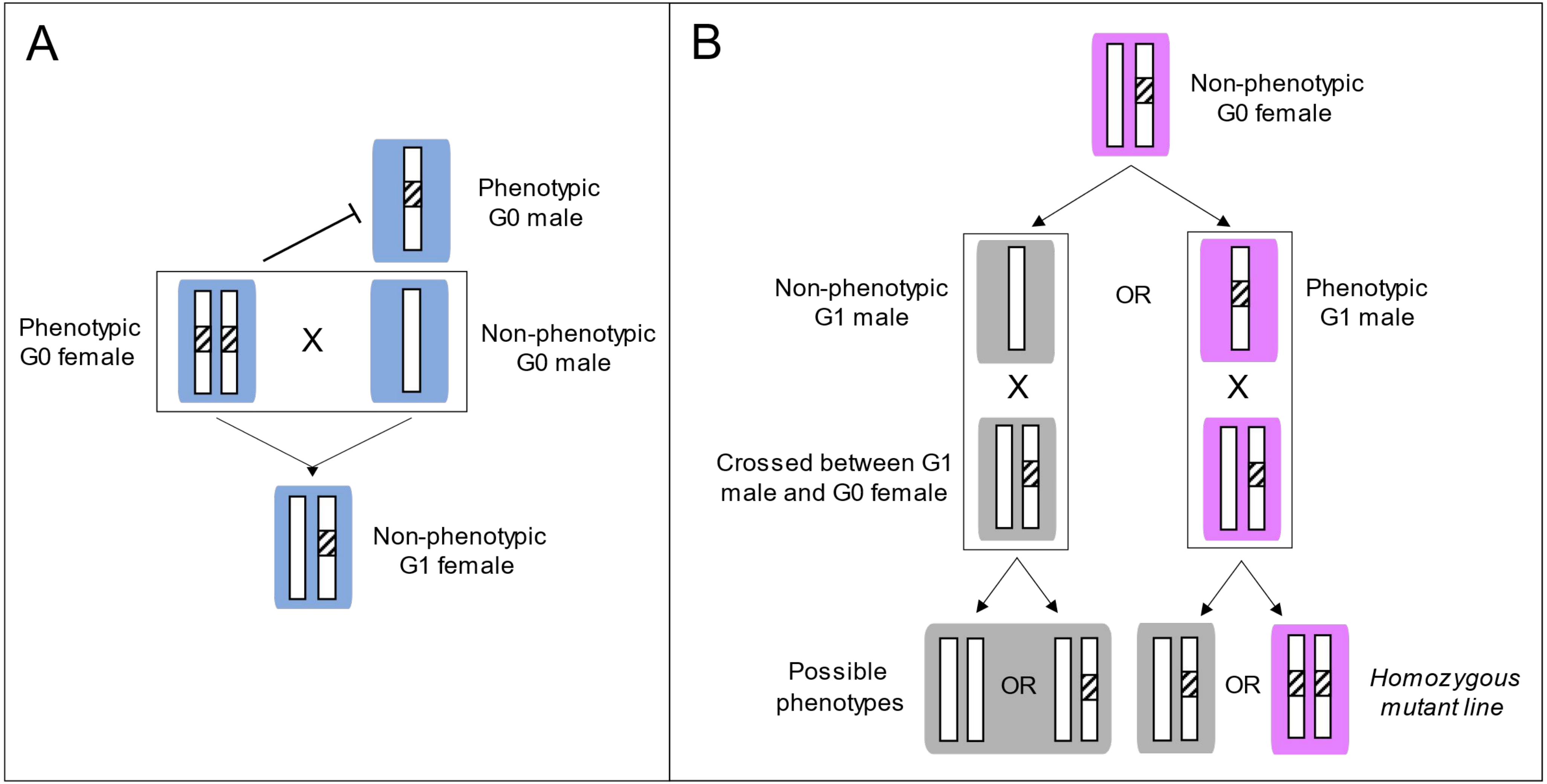


**Figure S2**. Genetic explanation for the failure and success of establishing *cinnabar* mutant line. A) Blue squares show the potential events occurred to produce non-phenotypic heterozygous progenies. Mating might take place between the phenotypic G0 female and an non-phenotypic G0 male before this female was collected for downstream screening. B) Pink squares represent the potential scenario of obtaining phenotypic homozygous mutant line from non-phenotypic G0 female. Grey squares represent events that did not occur or occurred but not chosen for downstream screening.
