## Supplementary Tables S1 & S2 for "CRISPR/Cas9-mediated genome editing of *Frankliniella occidentalis,* the western flower thrips, via embryonic microinjection"

**Table S1**. Small guide RNAs (sgRNAs) designed for CRISPR-mediated knockouts of *white* and *cinnabar* genes in *Frankliniella occidentalis*

| **sgRNA name** | **sgRNA sequence** | **Target region** | **NCBI accession for genomic scaffold** |
| --- | --- | --- | --- |
| Fo_White_Ex2 | 3’ GGUGAGGGUGUUGAGCAGCG 5’ | *White* exon 2 | NW_026272940.1 |
| Fo_White_Ex4 | 3’ GAACAGCGAGAAGACCUCGG 5’ | *White* exon 4 | NW_026270734.1 |
| Fo_Cinnabar_Ex1 | 5’ AUGCGCGCCAGGAUGAUCCA 3’ | *Cinnabar* exon 1 | NW_026268346.1 |
| Fo_Cinnabar_Ex4 | 5’ GGAGCUCAGGAUACCUCCAA 3’ | *Cinnabar* exon 4 |  |

**Table S2**. Primer pairs designed to detect the genomic sequence deletions in *white* and *cinnabar* knockout lines of *Frankliniella occidentalis*

| **Primer name** | **Primer sequence (5’ to 3’)** | **Length** | **Annealing temperature (°C)** | **Amplicon size (bp)** |
| --- | --- | --- | --- | --- |
| Fo_White F1 | GAGCGACGTGAACGTCTTC | 19 | 58 | 1032 |
| Fo_White R1 | CTTATTGCTCTGTATACGCACTGG | 24 | 57 |  |
| Fo_Cinnabar F1 | ACAGAGGTGCACATTTCCCA | 20 | 58 | 1003 |
| Fo_Cinnabar R1 | TCTGATAAATCTGCGCGTGG | 20 | 58 |  |
| Fo_Cinnabar F2 | CATCAACCTGGCCATGTCG | 19 | 58 | 770 |
| Fo_Cinnabar R2 | GTTCGGCAGGTAAAGCAAGT | 20 | 57 |  |
