## Supplementary File 1 Microinj Protocol for "CRISPR/Cas9-mediated genome editing of *Frankliniella occidentalis,* the western flower thrips, via embryonic microinjection"

**SUPPLEMENTARY FILE 1:** Microinjection protocol for CRISPR/Cas9 delivery into *Frankliniella occidentalis* embryos

prepared by: Lucas de Oliveira and edited by William Klobasa & Jinlong Han

**NOTE:** This detailed protocol was contributed as part of the paper entitled "CRISPR/Cas9-mediated genome editing of *Frankliniella occidentalis,* the western flower thrips, via embryonic microinjection" by Jinlong Han[1,2*], William Klobasa[1*], Lucas de Oliveira[1], Dorith Rotenberg[1], Anna E. Whitfield[1] and Marcé D. Lorenzen[1].

[1]Department of Entomology and Plant Pathology, North Carolina State University, Raleigh, NC 27695, USA

[2]Current address: Department of Agricultural Biology, Colorado State University, Fort Collins, CO 80523, USA

*These authors contributed equally to this work

The protocol described here was developed for *F. occidentalis* rearing methods and conditions developed by Ullman et. al., 1992.

### Materials and Tools:

- 1 L Erlenmeyer flask - 1 unit
- 100 mL sterile media bottle with a cap - 1 unit
- 140 mm diameter petri dish (Thermo Scientific™ Product code 11339283) - 1 unit
- 16 oz. deli cups - 3 cups
- 35mm diameter stackable petri dishes (Genesee Scientific Cat #: 32-103) - 4 to 5 dishes
- 70% ethanol - 1 spray bottle
- 90 mm diameter petri dish with lid (Thermo Scientific™, Cat. No. 101R20) - 4 dishes
- Agar - 1.44 g
- Ahlstrom filter paper (9cm diameter, black. Cat. 8613-0900) - 1 pack of 100 filters
- Bubble level - 1 unit
- Cafeteria tray - 1 unit
- Circular coffee filter - 1 pack containing 100 filters
- DI water - 244 mL
- Double-sided tape - 1 roll
- Eppendorf™ Femtotips™ Microloader P20 Tips for FemtoJet Microinjector (Fisher Catalog number: 930001007) - 1 box containing 96 tips
- Fine synthetic paintbrush (Winsor & Newton, Series 223, Size 000) - 1 unit
- Fresh green bean pods (Phaseolus vulgaris) - steady supply for maintaining the colonies
- Funnel - 1 unit
- Gene-specific sgRNA
- Green food coloring - 1 bottle
- Invitrogen™ TrueCut™ Cas9 Protein v2 (Thermofisher, Cat. No. A36499) - 1 tube
- Kimwipes - 1 box

- Large forceps - 1 pair
- Large soft brush (Winsor & Newton, Series 240, Size 2) - 1 unit
- Microscope cover glass (Fisher Scientific 12-540B, 22x22-2) - 4 to 5 coverslips
- Microscopy slide - 1 slide
- PCR strip tube - 1 tube
- Paper towel - 1 package
- Parafilm - 1 roll
- Quartz microcapillary tubes without filament (Sutter Instrument Company, O.D. = 1.0 mm, I.D. = 0.50 mm; 10 cm length; Item No. Q100-50-10) - 1 pack containing 100 tubes
- Razor blade - 1 blade
- Scissors - 1 pair
- Sucrose - 2.4 g
- Synthego EZ sgRNA kit – modified (This includes synthego 80-mer SpCas9 scaffold, as well as 2'-O-Methyl at 3 first and last bases, 3' phosphorothioate bonds between first 3 and last 2 bases) - 1 kit
- Ultra-fine forceps (Fisher Scientific model 12-000-122) - 1 pair
- Water squirt bottle - 1 unit
- Whatman paper (GE Healthcare Life Sciences, Grade 1, 85mm diameter) - 1 pack of 100 filters
- White paper - 2 sheets
- Wildtype thrips adults in colony cups - 2 to 4 cups

### Equipment:

- AirClean® Systems AC600 Series PCR Workstation (Model AC632LFUVC - 49536)
- Eppendorf FemtoJet microinjector (Model No. 5247 000.013)
- Hand Control for FemtoJet 4i/4x (Part No. 5252070011)
- Laser-based micropipette puller (Sutter Instrument Company, Model No. P-2000)
- Leica Mz6 Microscope fitted with with AmScope GT100 X-Y Gliding Table
- Micropipette beveler (Sutter Instrument Company, Model No. BV-10)
- Microwave
- Modular incubator chamber (Billups-Rothenberg Pat. No. 5352414)
- PCR strip Mini-centrifuge (Genemate Bioexpress Mini-centrifuge, Model C-1301)
- Refrigerator
- Temperature controlled incubation chamber (Percival Model AL22L2 Serial 25865.01.18)

### Terminology:

- **Oviposition dish:** This refers to the petri dish bottom filled with 3% sucrose solution, covered with a thin layer of stretched parafilm and fortified with additional parafilm along the sides.

- **Egg lay:** This refers to the three hour window in which the thrips are allowed to oviposit their eggs into the oviposition dish

- **Deli cup:** This refers to 16 oz deli containers, often used in the food service industry, and are used to hold and rear thrips.
- **Oviposition cup:** This refers to a deli cup that contains a petri dish cover housed within to serve as a shelf to set the oviposition dish upon.
- **Colony cup & colony cup lid:** This refers to a 16 oz deli cup used to rear and store thrips. The snap-on lids of the deli cup were modified by cutting a 7.5 cm hole and 178-mesh monoester silk screen (NBC Industries Co., Tokyo, Japan) was hot glued to the underside of the lid. This mesh prevents thrips from escaping while providing ventilation. Inside these deli cups, a 85mm diameter Whatman filter paper was placed at the bottom to absorb the moisture and green bean pods were used as a food source for the thrips.
- **Mini-centrifuge adapter:** This refers to a makeshift adapter used to spin loaded microinjection micropipettes in a centrifuge to eliminate air bubbles in the micropipette. To make the adapter, take a PCR tube and cut the bottom with scissors, then make a hole in the cap with a razor blade. Take a capillary tube and wrap it with parafilm until it is the same diameter as the PCR tube. Insert the parafilm with capillary tube into the modified PCR tube, then remove the capillary tube. The mini-centrifuge adapter can be reused.
- **Hatching dish:** This refers to a 35 mm petri dish used to house eyespot developed thrips embryos during their transition to first instar larvae. The hatching dish is used to maintain thrips hydration, prevent thrips escape, and minimize damage to thrips since embryos are easier to transfer than larvae. To prepare a hatching dish, place two circles of 30 mm Ahlstrom filter paper into a 35 mm diameter petri dish and spritz with water to moisten. Cut squares of Parafilm to fit over the lids of the hatching dish. Once thrips embryos are in the hatching dish, Place the Parafilm over the open dish and cover with the petri dish lid. Use additional Parafilm to seal the outside edge of the petri dish to prevent thrips escape.

### Methodology:

##### Part 1: Micropipette Pulling and Beveling

1. Adjust the settings of the Sutter P-2000 micropipette puller to the following parameters: heat = 700, filament = 004, velocity = 040, delay = 170, and pull = 160.
2. Insert a 10 cm quartz microcapillary tube (without filament) into the machine and initiate the pulling process.
3. Store the micropipette inside a 140 mm petri dish fitted with a slitted foam strip.
4. Repeat steps 2-3 until the desired number of micropipettes are produced.
5. Prepare a Sutter BV-10 micropipette beveler fitted with a fine quartz grinding plate assembly and an 80X optical attachment.
   *Note: For a detailed explanation of the grinding plate assembly process, see the Sutter BV-10 operation manual (Sutter 2018).

6. Raise the micropipette manipulator to its highest setting, then adjust the beveling angle to 25-30 degrees. Position the manipulator so the pipette tip can be lowered onto the beveling surface two-thirds from the center of rotation. Secure the micropipette in the clamp on the manipulator, tightening the nylon washer.
   *Note: Beveling on the outer edge or the center of the grinding plate should be avoided.

7. Turn on the BV-10 beveler.

8. Lower the micropipette close to the grinding plate using the coarse adjustment. Using the 80X optical attachment, align the micropipette until it is centered in view by adjusting the manipulator. Use the focus and fine height adjustment as needed. Position the light source to cast a shadow of the micropipette on the rotating grinding plate. With the fine adjustment, lower the micropipette onto the grinding plate until the tip of the micropipette and its shadow appear to meet. Carefully continue lowering the micropipette until a subtle flex can be seen.

9. Begin a 15 second mental count. Monitor if the micropipette retains its slight flex during this time. If the tension relaxes and the flex disappears, lower the micropipette slightly further with the fine adjustment until the slight flex is visible again. Also, monitor the tip; there should be a barely visible bevel on the micropipette as it is reaching completion. Once 15 seconds have elapsed, raise the micropipette away from the grinding plate using the fine adjustment, and then further with the coarse adjustment until the manipulator is at or near its highest position.
   *Note: Use your best judgment to determine if a micropipette was properly beveled. Poorly beveled micropipettes should be discarded in the appropriate waste container.

10. Release the micropipette from the manipulator and store it in a 140 mm petri dish fitted with a slitted foam strip.

11. Repeat steps 8-11 until the desired number of micropipettes have been beveled.

12. Label the micropipette container with the pull settings, if beveled or unbeveled, and the date.
    *Note: These micropipettes can be used long after they are pulled and beveled so long as they are free from contamination and dust. To avoid losing batches of micropipettes, minimize how long their container is open.

**Part 2: Oviposition Dish Preparation**

1. Prepare an 80 mL solution of 3% sucrose in DI water and add 40 uL of green food coloring. Store the solution in a 100 mL sterile media bottle with a cap.
   *Note: Solution can be prepared up to 2 weeks in advance and stored in a refrigerator, but fresh preparation is recommended. Inspect for visible signs of microbes before use.

2. Cut a sheet of parafilm approximately 7 cm x 5 cm. Stretch the parafilm lengthwise and widthwise until it is as thin as possible to allow an "easy" puncture by the thrips ovipositor. Ensure the parafilm sheet is at least 10.5 cm x 10.5 cm to fit over the petri dish.

3. Cover the bottom housing of a 90 mm diameter petri dish with the stretched parafilm, leaving an opening along the rim approximately 0.5 cm wide. Ensure the parafilm is secured to the sides of the petri dish.

4. Hold the petri dish at an angle and position the opening to be at the highest point. Fill the dish through the opening at the top until the petri dish is full (Figure 1A - 1B). As the dish reaches its maximum capacity, the solution will cause a bulge in the parafilm.

5. Grasp the Parafilm where the opening is. To seal the oviposition dish while avoiding the presence of air bubbles, begin tilting the petri dish back into a level position. Simultaneously, pull the parafilm to cover the opening (Figure 1C - 1E).

6. Trim off the excess parafilm of the top sheet against the bottom edge of the petri dish.

7. Cut a strip of parafilm approximately 1.5 cm x 10 cm and wrap the outer perimeter of the petri dish with the parafilm strip. This will be referred to as the oviposition dish for the remainder of the protocol.
   *Note: To fortify the structural integrity of the top sheet of parafilm, when wrapping the perimeter of the dish, ensure approximately 0.5 mm of the parafilm strip extends over the side of the dish onto the top sheet of parafilm.

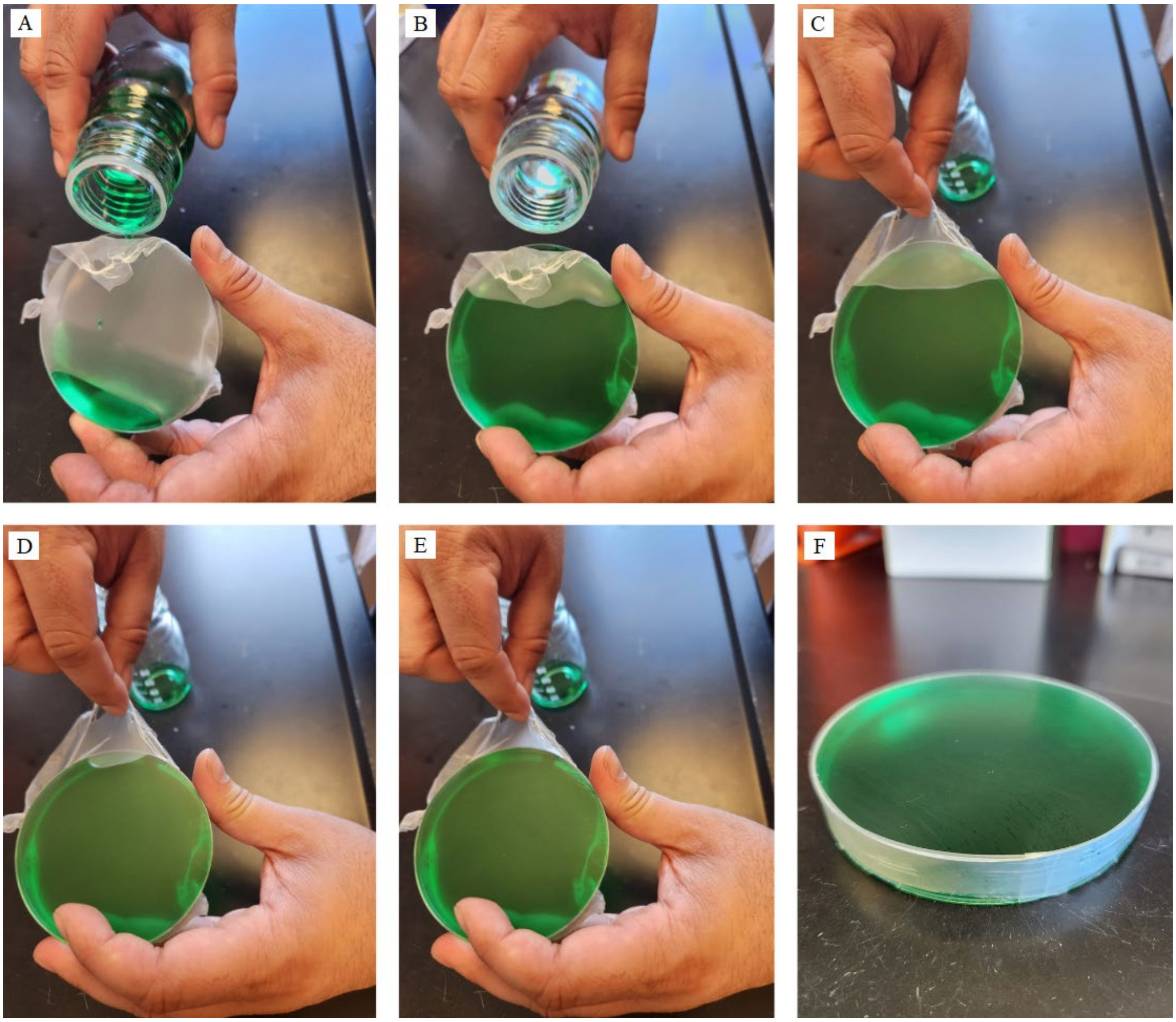

**Figure 1:** Oviposition dish construction. (A) - (E) showcases the process of filling and sealing the oviposition dish without introducing air bubbles. (F) depicts a completed oviposition dish with the perimeter wrapped in parafilm.

**Part 3: Egg Lay Setup**

1. Retrieve a clean deli cup and colony cup lid as well as a 90 mm diameter petri dish lid.

2. Cut a strip of parafilm and wrap the perimeter of the petri dish cover. Place the petri dish lid into the deli cup with the top of the lid facing down, ensuring that it is level when nested within the deli cup.

3. Press the petri dish lid into the deli cup, and the parafilm along its edge will make a seal. *Note: This will secure the petri dish lid in place within the deli cup, prevent thrips from escaping into the bottom chamber of the deli cup, and serve as a shelf to set the oviposition dish upon. This deli cup will be referred to moving forward as the oviposition cup.

4. Take two Kimwipes and place them on a flat surface overlapping in a perpendicular fashion then place the oviposition dish in the center of the Kimwipes (Figure 2A).

5. Using the Kimwipes as a sling, lower the oviposition dish into the oviposition cup. Ensure that the oviposition dish is level when nested within the oviposition cup. Set aside the oviposition cup.

6. Retrieve three to four colony cups containing wildtype thrips adults that are 2 - 12 days post eclosion from pupae.
*Note: Under our colony maintenance conditions, wildtype colony thrips were maintained at 23- 25 °C and we estimate under these conditions from egg to adult eclosion takes 11-12 days. WFT females have a preoviposition period of 1-2 days but remain reproductively efficient for the next 9-10 days (Kumm 2002). Theoretically, under our conditions, female thrips ages 2-12 days post eclosion from pupae should be viable for use in the egg lay protocol. However, we did not use adults that were younger than two days or older than seven days post eclosion from pupae, as we noticed reduced embryo yield and viability.

7. Open the lid of the colony cup and clean the thrips from the green beans and Whatman paper within. Place the beans and Whatman paper into a clean and empty deli cup. To clean off the thrips, grasp the beans with large forceps and tap the forceps with a metal implement to remove the thrips.
*Note: We used medium sized scissors as the tapping implement; however, the particular implement doesn't matter so long as it is metal and has sufficient mass.

8. By this point, the colony cups should only contain thrips and nothing else. Tap the bottom edge of the colony cup on the bench until the majority of the thrips are concentrated within a small area.

9. Retrieve the oviposition cup and place it on the bench. Grab the colony cup and begin to pour the thrips into the oviposition cup. Using a metal implement of choice, tap the colony cup to get more thrips into the oviposition cup.
*Note: It is not necessary to transfer all of the thrips from the colony cup into the oviposition cup. Doing so may risk the thrips becoming stressed due to overhandling.

10. Once the majority of the thrips have been transferred to the oviposition cup, place the colony cup on a flat surface. Retrieve the deli cup containing the beans and Whatman paper originally within the colony cup. Return the Whatman paper followed by the beans to the colony cup. Close the colony cup with its lid and set aside.

11. Repeat steps 8-12 for the other retrieved colony cups.

12. Seal the oviposition cup with a clean colony cup lid.
*Note: See terminology section for instructions on how to prepare a colony cup lid.

13. Place the oviposition cup within a temperature controlled incubation chamber (Percival Model AL22L2 Serial 25865.01.18) with the device set to 25°C with a 16 hour photoperiod.

14. Cover the oviposition cup with a filter paper to block direct light and allow 3 hours for the egg lay to be complete.

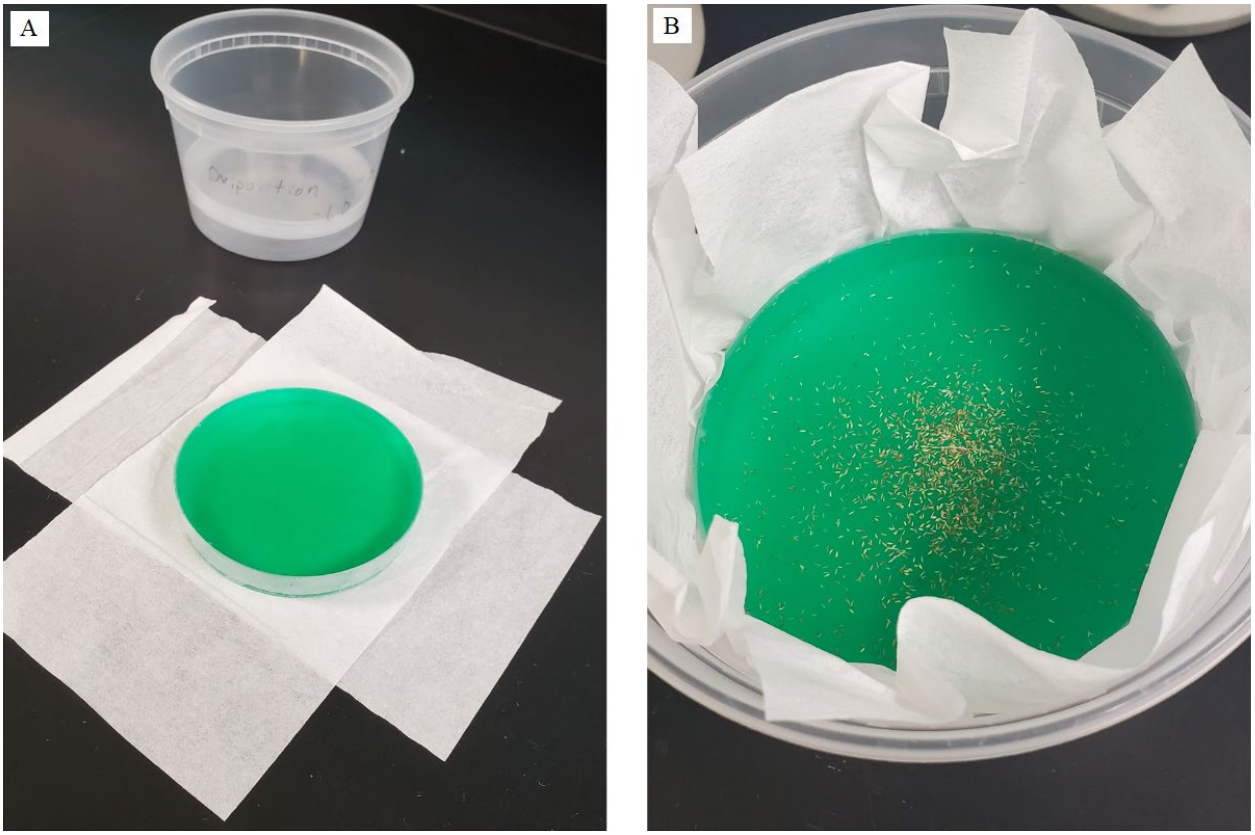


**Figure 2:** Egg lay. (A) depicts the oviposition dish on top of the Kimwipe sling in the foreground and the oviposition cup in the background. (B) depicts the oviposition dish after it has been lowered into the oviposition cup and thrips have been transferred to the surface of the oviposition dish.

**Part 4: Agar petri dish preparation**

*Note: This part of the protocol can be conducted while waiting for the egg lay. These agar dishes will maintain thrips' hydration levels later during microinjection.

1. In an 1 L Erlenmeyer flask, mix 1.44 g of agar with 144 mL of DI water.
2. Microwave the solution until the agar is completely dissolved.
3. Retrieve three 90 mm diameter petri dishes and find a level surface. You can verify the evenness of the surface with a bubble level. Place one of the petri dishes onto the level surface.
4. Wait for the agar to cool then pour the agar solution into the Petri dish until it is full to the brim.
   *Note: This agar dish will be used to hold microscopy coverslips containing thrips embryos on top of the microscope stage during the microinjection phase of the protocol.

5. Place the other 2 petri dishes on the bench and pour the remaining agar in roughly equal amounts between the two remaining petri dishes.
   *Note: It is not necessary to ensure that the other two petri dishes are on a level surface, only the single dish that was filled to the brim. One of these agar petri dishes will be used later in the protocol to hold microscopy coverslips containing thrips embryos while awaiting injection. The other will be used to incubate the thrips post-injection.

6. Allow the agar to solidify completely before use.

**Part 5: Egg Collection**

*Note: During the later microinjection phase of the protocol, you will notice embryos will become unreceptive to the delivered treatment solution (approximately 4-6 hours post oviposition). We hypothesize that at this point, the embryos are beginning to cellularize and reach the blastoderm stage, as it is known that WFT embryos reach this state within the first couple hours after oviposition (Kumm 2002). Therefore, the remainder of the protocol must be completed with this time limit in mind.

1. Place sheets of white paper on a benchtop to provide contrast for spotting thrips and ease of retrieval.

2. Retrieve the oviposition cup and the colony cups that originally housed the thrips. Open the lids and place them on the bench.

3. Grasp the Kimwipe sling and carefully raise the oviposition dish out of the oviposition cup. Place the oviposition dish and Kimwipe sling on a sheet of paper.

4. Carefully lift the oviposition dish out of the Kimwipe sling and place it elsewhere on a sheet of paper.

5. Take the first Kimwipe and return the thrips on it to one of the colony cups. Repeat this for the other Kimwipe.
   *Note: Ensure that roughly an equal number of thrips are returned to each colony cup.

6. Retrieve a large soft brush and wipe off any thrips on the oviposition dish into the colony cups.

7. Retrieve the oviposition cup and use the brush to carefully move any remaining thrips into the colony cups.

8. Bend the sheets of paper to concentrate any thrips into a line at the center of the sheet, then pour them into the colony cups.

9. Inspect the worktop surface for any thrips and use the large soft brush to sweep them into the colony cups.

10. Seal the colony cups with their lids and return them to their storage location.

11. Using 70% ethanol, spray and wipe down the worktop surface to kill any possible remaining thrips.

12. Retrieve a large Erlenmeyer flask, a funnel, a 140 mm diameter petri dish and a circular coffee filter.

13. Fold the coffee filter in half then fold it in half again. Place the tip of the folded coffee filter at the center of the 140 mm diameter petri dish and trace the coffee filter along where the outer perimeter of the petri dish is. Cut the coffee filter along the traced line with scissors.

14. Open up the folded filter to make a cone shape and place it inside the funnel. Then place the funnel into the opening of the Erlynmeyer flask and wet the filter paper with water.

15. Scrape the rim of the oviposition dish to cut open the parafilm top sheet. Grab the parafilm sheet and lift it up. Begin pouring out the contents of the oviposition dish into the filter. Once the dish is empty, remove the rest of the parafilm sheet (Figure 3A - 3D).

16. Rinse out the oviposition dish thoroughly with water from a squirt bottle to ensure all eggs are released from the dish (Figure 3E - 3F).

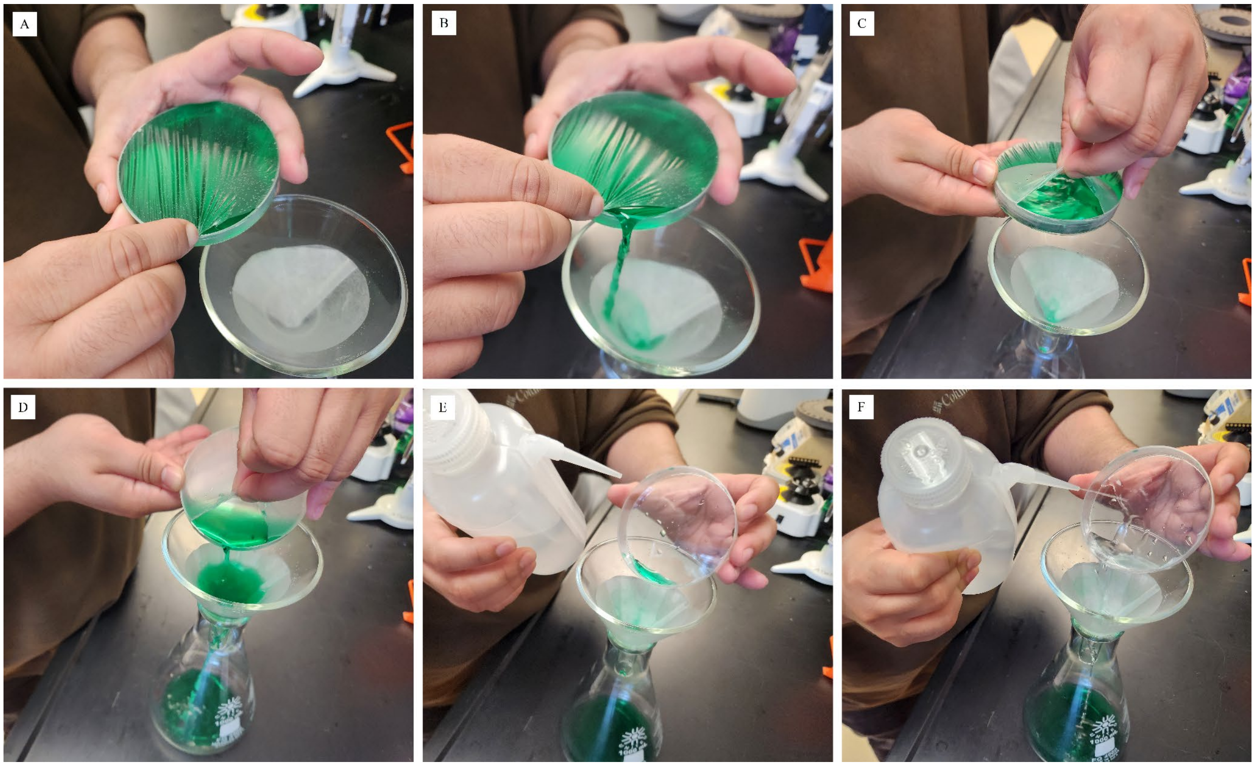


**Figure 3:** Egg Collection technique. (A) - (B) display how to properly open the oviposition dish. (C) - (D) show how to manipulate the parafilm sheet while draining the dish. (E) - (F) depict the wash step.

**Part 6: Slide Preparation**

1. Affix double-sided tape to a microscopy slide. Press a razor blade onto the tape to cut a thin sliver approximately 17 mm x 0.5 mm in size.

2. Peel the tape sliver from the microscopy slide and carefully affix it to a microscopy coverslip. Repeat this step for the number of cover slips needed, which should be equal to the number of embryos intended for injection / 20.
   *Note: It may be helpful to use a razor blade to lift up one end of the tape strip then use forceps to peel it off the microscopy slide. This minimizes marring the tape which could introduce peaks and valleys that make microinjection more difficult.

3. Under the microscope, use a type 000 fine synthetic paintbrush to transfer embryos onto the microscopy slide. Ensure that the posterior pole of the embryos is oriented towards the point where the micropipette will penetrate.

4. Line up 20 embryos per slide, then use forceps to transfer the slide to one of the two half filled agar petri dishes. Cover to maintain hydration.
   *Note: Five microscopy slides fit per agar petri dish.

5. Repeat step 4 until a desired number of embryos are plated onto the coverslips.
   *Note: When deciding how many embryos inject, keep in mind the time constraints due to cellularization. We recommend injecting 100 embryos per microinjection session, which would require 5 cover slips to be plated with embryos.

**Part 7: Micropipette Preparation**

1. Dilute Synthego sgRNAs to 400 ng/ul and dilute Invitrogen Truecut Cas9 protein v2 to 714 ng/ul in a 3.5ul total volume. Gently mix the components then briefly centrifuge. For proper complexing, incubate the tube at room temperature for 15 minutes then store the mix on ice.

2. Fill a pre-pulled micropipette with approximately 1 uL of the CRISPR/Cas9 and guide solution using the P20 Microloader Tips designed for the FemtoJet Microinjector.
   *Note: The Microloader tips can be labeled and reused if more injections will be performed with the same CRISPR/Cas9 solution in future.

3. Place the microloader tip into the open end of the micropipette. Depress the pipette and pull out quickly at the same time.
   *Note: If frequent micropipette clogging is occurring later in the protocol, try loading micropipettes under a PCR workstation hood to prevent contamination.

4. If any air bubbles form, load the micropipette into the mini-centrifuge adapter (see Terminology), then load into a PCR strip mini-centrifuge and run for 1 second. Repeat if air bubbles persist.

5. Load the micropipette into the FemtoJet. To do so, unscrew the universal capillary holder at the tip of the microinjector and insert the glass micropipette until the open end of the micropipette is flush with the back of the holder. Then insert the glass micropipette along with the holder into the microinjector and screw in to secure it.
   *Note: It is recommended to use the Eppendorf hand control accessory for the FemtoJet to easily dispense injections.

6. Turn on the Eppendorf FemtoJet microinjector. Allow time for the device to pressurize.

7. Set the device to the following parameters: Injection pressure 16 psi, Injection time 0.2 seconds, Holding pressure 5 psi
8. Press inject. The treatment solution should accumulate as a bead on the tip of the micropipette. If the liquid flows back up the micropipette, attempt to wipe the tip of the micropipette with a fine tipped brush wetted with water to disrupt the adhesive force of water.
9. If the issue persists, there could be dust within the tip restricting the flow of liquid. If dust is visible within the micropipette, repeat the procedure from step 1 with a new micropipette.

**Part 8: Microinjection**

1. Retrieve a single microscopy slide with embryos from the half filled agar dish and transfer it to the level, brim-filled agar dish.
2. Place the brim-filled agar dish with the embryo coverslip under a microscope on an x- and y-axis travel sliding stage and carefully position the injector such that the micropipette is on top of the microslide.
3. Using the “Empty” function in the Control Station, apply pressure for 2-3 seconds.
4. Adjust the height and position of the micropipette until it is poised to inject the first embryo. Then inject CRISPR/Cas9 solution into the first embryo, being sure not to inject too deeply. Retract the micropipette and adjust the x- and y-axis travel sliding stage so that the next embryo is in position.
   *Note: If the micropipette clogs, use a fine tip brush to dislodge the debris. If it cannot be unclogged, prepare a new micropipette following the instructions in Part 7.
5. Repeat step 4 for all 20 embryos on the microscopy coverslip, then transfer the coverslip with the injected embryos to the second half-filled agar petri dish and cover it.
6. Move the half-filled agar petri to the modular incubator chamber (Billups-Rothenberg Pat. No. 5352414) with a single damp paper towel on the bottom of the chamber. Then place the chamber into a temperature-controlled incubation chamber (Percival Model AL22L2 Serial 25865.01.18) at 25°C with a 16-hour photoperiod.
7. Repeat steps 1-6 until all embryos are injected.
8. Incubate embryos for 62 hours.

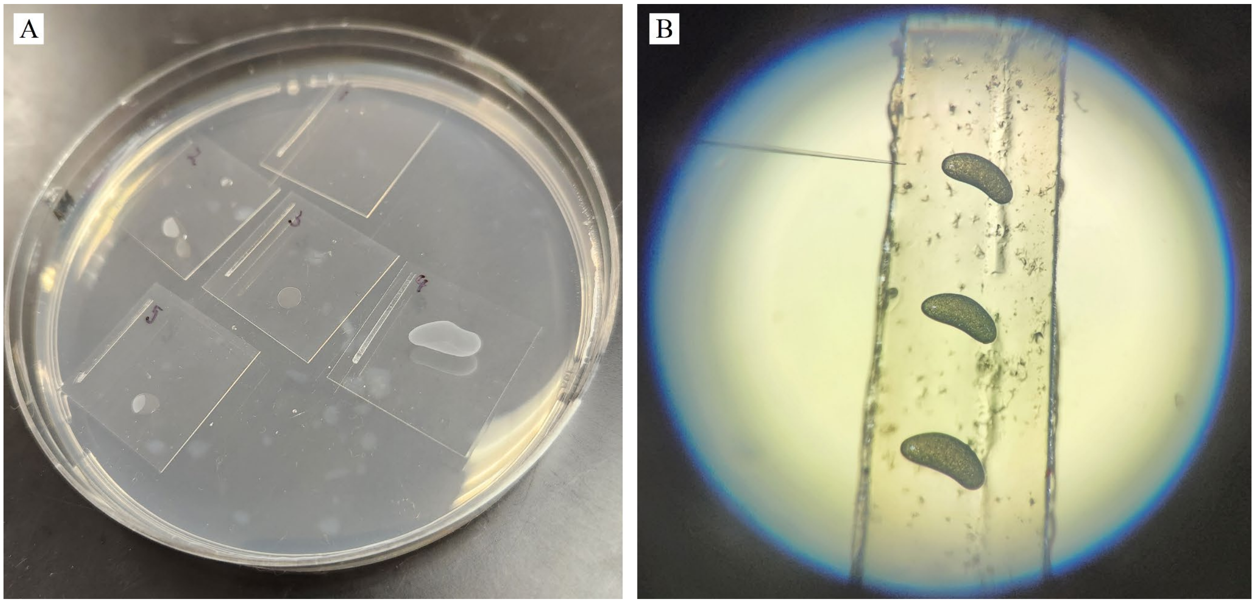


**Figure 4:** (A) depicts one of the half-filled agar petri dishes containing embryos ready for injection. The embryos are mounted on microscopy coverslips with a strip of double sided tape. (B) shows embryos on the coverslips under the microscope with the micropipette positioned for injection.

**Part 9: Hatching Dish Transfer**

1. Prepare a hatching dish by taking 35 mm diameter petri dishes and black Ahlstrom filter paper. Cut circles of the black Ahlstrom paper to fit inside the petri dish, and insert 2 circles into the petri dish. Spritz with water to moisten them.

2. Cut squares of Parafilm to fit over the lids of the hatching dish.

3. Examine the slides with injected thrips under a microscope. If eyespots can be clearly seen, transfer them into a hatching dish using a fine tip brush and water.

4. Place the Parafilm over the open dish and cover with the petri dish lid. Use additional Parafilm to seal the outside edge of the petri dish to prevent thrips escape.

5. Repeat steps 1-4 until all eyespot developed embryos have been moved to hatching dishes.

6. Transfer the hatching dishes back to the modular incubator chamber with a single damp paper towel on the bottom of the chamber. Place the modular incubator chamber within a temperature controlled incubation chamber at 25°C with a 16 hour photoperiod for 24 hours.

**Part 10: Colony Cup Creation & Maintenance**

1. Obtain a deli cup, 3 fresh green beans, an 85mm Whatman filter paper, and a clean colony cup lid. Place the Whatman paper in the deli cup then add the 3 green beans.

2. After the 24 hour incubation, examine thrips within the hatching dishes. Using a fine tip brush and water, transfer L1 thrips from the hatching dish onto the green beans within the deli cup.

3. Secure the lid onto the deli cup, cut two 10 cm x 2 cm parafilm strips, and double wrap the rim of the cup with the parafilm.
*Note: It is important to seal mutant colony cups with parafilm to prevent colony contamination. The colony cups themselves are not secure and thrips are able to enter and exit from the gap between the deli cup and lid.

4. Store the newly created mutant colony cup within a temperature controlled incubation chamber at 25°C with a 16 hour photoperiod.

5. To maintain the mutant colony, provide fresh green beans to the deli cup every 3 to 4 days and remove any dried or moldy beans to prevent contamination and maintain a healthy food source for the thrips.
*Note: Be sure not to overcrowd the container as this can lead to moisture build-up and mold growth. Filling a colony cup with more than 6 green beans often results in these issues.

6. Use the mutant thrips colony for future experiments as needed.
